# Structural comparisons of human and mouse fungiform taste buds

**DOI:** 10.1101/2024.07.10.602971

**Authors:** Brigit High, Thomas E. Finger

**Author notes:** Corresponding Author: Brigit High, Dept. Cell & Devel. Biology, Univ. Colorado School of Medicine MS 8108. Room L18-11118, RC-1 12801 E. 17th Ave., Aurora CO 80045, Voice.

## Abstract

Taste buds are commonly studied in rodent models, but some differences exist between mice and humans in terms of gustatory mechanisms and sensitivities. Whether these functional differences are reflected in structural differences between species is unclear. Using immunofluorescent image stacks, we compared morphological and molecular characteristics of mouse and human fungiform taste buds. The results suggest that while the general features of fungiform taste buds are similar between mice and humans, several characteristics differ significantly. Human taste buds are larger and taller than those of mice, yet they contain similar numbers of taste cells. Taste buds in humans are more heavily innervated by gustatory nerve fibers expressing the purinergic receptor P2X3 showing a 40% higher innervation density than in mice. Like Type II cells of mice, a subset (about 30%) of cells in human taste buds is immunoreactive for PLCβ2. These PLCβ2-immunoreactive cells display CALHM1-immunoreactive puncta closely apposed to gustatory nerve fibers suggestive of channel-type synapses described in mice. These puncta, used as a measure of synaptic contact, are however significantly larger in humans compared to mice. Altogether these findings suggest that while many similarities exist in the structural organization of murine and human fungiform taste buds, significant differences do exist in taste bud size, innervation density, and size of synaptic contacts that may impact gustatory signal transmission.

## Introduction

The morphology of mammalian taste buds, including mice and humans, is largely conserved across species. Both mouse and human tongues contain fungiform papillae with taste buds innervated by gustatory afferents arising from the chorda tympani nerve (Arvidson and Friberg 1980; Cheng and Robinson 1991). These nerve fibers receive synapses from the taste receptor cells and express purinergic receptors required for successful gustatory transmission (Finger *et al*. 2005). However, P2X receptor subunit composition differs between the two species (High *et al*. 2023). We therefore hypothesize that while major features of fungiform taste buds may be preserved between species, other features may diverge.

Rodent fungiform taste buds, which reside within the anterior two-thirds of the tongue, have been extensively studied in terms of morphological and molecular features while those of humans have not been characterized in as much detail. Moreover, mice are commonly used as a model organism for human taste, but it is unclear to what degree mouse and human taste buds are similar. Prior work describes shared structural, morphological, and molecular features in circumvallate and laryngeal taste buds of mice and humans (Jette *et al*. 2020; Tizzano *et al*. 2015), however, some differences are known to exist in the distribution of fungiform taste buds in humans compared to mice. A mouse typically has between 90-110 fungiform papilla each containing a single, taste bud shaped like a garlic (Guagliardo and Hill 2007; Liu *et al*. 2009). In contrast, humans contain approximately 200 fungiform papillae with between 0-15 taste buds per papillae, with an average of 3.6 taste buds per papilla in the 60-70% of papillae that do contain taste buds (Cheng and Robinson 1991; Miller 1989; Miller and Reedy 1990).

While gross anatomical differences between mice and humans are documented in the distribution and number of fungiform papilla, morphological and molecular details are less well-characterized. For example, several key proteins required for type II taste cell transduction and transmission have been identified in both mouse and human taste buds via immunohistochemistry. Taste cells of mouse and human circumvallate taste buds are immunoreactive for Type II cell transduction cascade elements, PLCβ2 and GNAT3; intragemmal fibers in both species are immunoreactive for the purinergic receptor P2X3 (Tizzano *et al*. 2015). Such work suggests that requisite cellular constituents of type II signaling mechanisms are conserved between rodents and humans, and that purinergic neurotransmission is an essential feature of taste buds in both mouse and humans. Although both mouse and human taste buds employ purinergic neurotransmission, details of their systems differ. Our recent work reports that P2X2, which is largely co-expressed in mouse gustatory afferents along with its binding partner P2X3, is infrequently detected in humans (High *et al*. 2023). This begs the question of what other differences may exist in gustatory signal transmission between mice and humans. For example, in mice, type II taste cells form channel-type synapses with gustatory afferents where ATP is released directly into the synaptic cleft from the taste cell through the large-pore CALHM1 channel, which can be used as a marker for synaptic contact (Kashio *et al*. 2019; Romanov *et al*. 2018; Taruno *et al*. 2013). Whether human taste buds share this feature is unknown. Here, we describe findings comparing morphological and molecular features related to innervation and synaptic contact present in mouse and human taste buds.

## Methods

### Human tissue

Human adult fungiform taste buds were obtained from 13 subjects (6 females, 7 males, aged 22-33 years) old at the University of Colorado Hospital Outpatient Clinical and Translational Research Center. These subjects gave informed consent and agreed to undergo fungiform papillary biopsies in which three fungiform papillae were removed from the anterior one-third of the tongue bilaterally without anesthesia using sterile iridectomy scissors (protocol 14-0439 approved by the Colorado Institutional Review Board). Demographic features of the subjects are in **Table 1**. All subjects were considered to be healthy with no known diagnosed diseases or illnesses or use of tobacco products in the last year.

**Table 1.** Demographics for human samples.

| **Subject** | **Age**<br>**(years)** | **Sex** | **Race/Ethnicity** | **# Taste buds evaluated** |
| --- | --- | --- | --- | --- |
| 1 | 22 | F | White, non-Hispanic | 3 |
| 2 | 26 | M | White, non-Hispanic | 1 |
| 3 | 24 | F | White, non-Hispanic | 1 |
| 4 | 25 | F | Native American, non-Hispanic | 2 |
| 5 | Adult | M | White, non-Hispanic | 1 |
| 6 | 30 | F | White, non-Hispanic | 2 |
| 7 | Adult | M | Unknown | 1 |
| 8 | 32 | F | Black, non-Hispanic | 3 |
| 9 | 31 | F | Black, non-Hispanic | 3 |
| 10 | 33 | M | White, non-Hispanic | 1 |
| 11 | 27 | M | White, non-Hispanic | 1 |
| 12 | 33 | M | White, non-Hispanic | 2 |
| 13 | 27 | M | White, non-Hispanic | 1 |
| **Summary demographics** |  |  |  |  |
| **# Subjects** | **Age range** | **Female vs.**<br>**male** | **Race/Ethnicity** | **Total # taste buds** |
| 13 | 22-33 yr | 6 female<br>7 male | 9 White, non-Hispanic<br>2 Black, non-Hispanic<br>1 Native American, non-Hispanic | 22<br>(14 female, 8 male) |

### Mouse tissue

Tongues from 7 C57BL/6J mice (4 females, 3 males, aged 3-10 months) were obtained with approval of the Animal Care and Use Committee at the University of Colorado Medical School. These tissues were immersion-fixed for 4 hours in 4% paraformaldehyde to mimic fixative conditions used for the human-sourced tissue.

### Immunohistochemistry and reagent validation

Antibodies in this study have been validated and used previously in other studies. Antisera against P2X3 have been tested in knockout mice and show staining consistent with other P2X3 antibodies (Finger *et al*. 2005). The GNAT3 antibody was validated with a 27kDA band in Western blots of human colon and testis lysates which was blocked successfully with an immunizing peptide (Aviva Systems Biology Cat# OAEB00418, RRID: AB_10882823). It was also used in prior immunohistochemical studies of taste buds both in mice and humans (Larson *et al*. 2015; Tizzano *et al*. 2015). CALHM1 antisera recognizes a 46kDA band in Western blots in HepG2 cell lysate (Kashio *et al*. 2019). This antibody was validated using a cognate peptide block as well and has been used in prior immunohistochemical studies in mice (Kashio *et al*. 2019; Romanov *et al*. 2018). Finally, PLCβ2 has been validated with cognate peptide blocks (60µM EPLVSKADTQESRL) and reacts with a 28kDA band on a Western blot of mouse brain (Thomas Finger - University of Colorado Cat# PLCβ2 Green, RRID: AB_2910247).

All human and mouse tissues were immersion-fixed for 4 hours using 4% paraformaldehyde (PFA) in phosphate buffered saline and cryoprotected in 20% sucrose in 0.1M phosphate buffer, pH 7.4 for three days at 4° C. The tissues were sectioned at 14µm-16µm on a cryostat and mounted directly on Superfrost Plus slides (Fisher Scientific) in a 1:3 or 1:4 series. After drying, slides were rinsed in deionized water, and then underwent antigen retrieval in buffer (1X Tris-EDTA, pH 9.0 for slides stained with GNAT3 or in 1X Tris-EDTA, pH 6.0 for slides stained with CALHM1) at a temperature of 85°C for 10 minutes. After being allowed to cool, the slides were rinsed three times for 5 minutes each in 0.1M phosphate-buffered saline, non-specific binding was blocked for at least 1hr at room temperature in blocking solution (2% normal goat serum, 1% bovine serum albumin, 0.3% Triton in PBS). Mouse slides stained with CALHM1 were also pre-incubated in unlabeled goat-anti-mouse IgG Fab fragment (Jackson ImmunoResearch, lot 152150) at 1:25 dilution for 2 hours at room temperature to reduce non-specific binding of secondary antibodies. All tissues underwent incubation with primary antibodies diluted in blocking solution. All slides were incubated with P2X3 as well as GNAT3 or CALHM1 primary antibodies. Slides with antisera for GNAT3 were incubated for two nights at 4°C while those with antisera against CALHM1 were incubated for 4-5 nights per the protocol in Romanov et al., 2019 (**Table 2**). After incubation with the primary antibodies, the tissue samples were rinsed with 0.1M PBS, pH 7.4 three times for 10 minutes per rinse. They were then incubated for 2 hours with fluorescent secondary antibodies at 1:800 dilution for all antibodies except CALHM1 (**Table 2**). CALHM1 was incubated with an isotype-specific secondary, against IgG2A (**Table 2**). Finally, the slides were washed twice for 10 minutes each in 0.1M PBS and one time for 10 minutes in 0.05M PB before being coverslipped with DAPI Fluormount (SouthernBiotech – Birmingham, AL, USA).

**Table 2.** Primary and secondary antisera.

| **Primary antisera** | **Marker for:** | **Company, Catalog No.** | **Research Resource Identifier No. (RRID)** | **Host; dilution** |
| --- | --- | --- | --- | --- |
| P2X3 | ATP receptor on taste nerves and airway afferents | Alomone, APR-016 | AB_2313760 | Rabbit; 1:500 |
| GNAT3 (α-gustducin) | G-protein subunit in Type II taste cells | Aviva Systems Biology, OAEB00418 | AB_10882823 | Goat; 1:500 |
| CALHM1 | Voltage-gated, ATP-permeable channel present on Type II taste cells | Philippe Marambaud, Feinstein Institute, CALHM1-32C2 | AB_2927681 | Mouse, 1:50 |
| PLCβ2 | Type II taste cells | Thomas Finger - University of Colorado, PLCβ2 Green | AB_2910247 | Guinea pig; 1:500 |
| **Secondary antisera** | **Host** | **Company, Catalog No.** | **Research Resource Identifier No. (RRID)** | **Host; dilution** |
| A488 | Donkey anti-rabbit | Invitrogen by ThermoFisher Scientific, A-21206 | AB_2535792 | 1:800 |
| A568 | Donkey anti-rabbit | Invitrogen by ThermoFisher Scientific, A-10042 | AB_2534017 | 1:800 |
| A568 | Goat anti-mouse IgG2A | Invitrogen by ThermoFisher Scientific, A-21134 | AB_2535773 | 1:500 |
| A594 | Donkey anti-guinea pig (IgG, H+L) | Jackson ImmunoResearch | AB_2340474 | 1:800 |
| A647 | Donkey anti-goat | Invitrogen by ThermoFisher Scientific, A-21447 | AB_141844 | 1:800 |
| A647 | Donkey anti-guinea pig | Invitrogen by ThermoFisher Scientific, A-21450 | AB_2735091 | 1:800 |

### Image analysis

Images of mouse and human taste buds were taken using a 63x (n.A. 1.4) objective on a Leica SP8 confocal microscope. Only longitudinally sectioned taste buds with visible taste pores were selected for further analysis as these represent sections through the centerline of the taste bud. All taste buds were preprocessed in ImageJ (Fiji) 1.53v and analyzed at a resolution of 1024×1024 pixels (185×185μm) in Imaris. Preprocessing included selecting a 6μm z-stack of images, followed by despeckling and subtracting background at a 50px (9μm) rolling ball radius. This function eliminated fine-grain non-specific labeling especially generated by the anti-mouse secondary in mouse tissues. Two-dimensional features - height, width, and taste bud longitudinal sectional area – were calculated in Fiji using the sample volume which was the centermost portion of the image stack, i.e. the 6 µm z-stack of images from the middle of the depth of the tissue section.

Methods for calculating the volume of the taste bud within the 6μm depth – referred to as the taste bud sectional volume – and innervation volume of the taste bud are based on those described in Ohman and Krimm, 2021. Imaris version 9.9.1 was used to calculate three-dimensional features, such as the taste bud sectional volume and innervation density. Measurements related to synaptic contact were also calculated within Imaris. Detailed instructions for how these metrics were determined are in **Supplementary methods**.

### Statistical analysis of immunofluorescent images

Statistical analysis was performed using GraphPad Prism. Unpaired t-tests were used to compare a variety of measurements taken in image stacks of mouse and human taste buds. This is also indicated in **Table 3**. Simple linear regression was used in measurements examining relationships between measurements. An Abercrombie correction was used to correct for bias in counting differently sized objects, as the width of mouse and human taste cell nuclei were found to be significantly different, and the corrected values are used to estimate the number of taste cells present within the sample (Abercrombie 1946; Coggeshall 1992; von Bartheld 2018). These values are used as a surrogate for the number of taste cell nuclear profiles counted in a representative 6μm image stack. In counting PLCβ2-positive taste cells, only immunoreactive profiles containing a nucleus within the sample volume were considered.

**Table 3.** Summary of results.

|  |  |  |  |
| --- | --- | --- | --- |
| *Unpaired t-test used in all calculations* | **Mouse (n=16)** | **Human (n=18)** | **p-value** |
| **Height (μm)** | 43.13±8.53 | 67.72±17.26 | p<0.0001* |
| **Width (μm)** | 34.55±8.71 | 32.30±9.35 | p=0.4723 |
| **Taste bud longitudinal sectional area (μm<sup>2</sup>)** | 1224±378.4 | 2054±888.4 | p=0.0014* |
|  | **Mouse (n=15)** | **Human (n=15)** | **p-value** |
| **Nuclear profile width (μm)** | 5.52±0.87 | 6.32±0.99 | p=0.0254* |
|  | **Mouse (n=16)** | **Human (n=12)** | **p-value** |
| **Taste cell count (raw)** | 19.81±6.03 | 21.08±6.37 | p=0.5945 |
| **Taste cell count (Abercrombie-adjusted)** | 10.33±3.14 | 10.27±3.10 | p=0.5945 |
| **Innervation density (%) \*** | 8.198±2.05 | 11.62±2.29 | p=0.0005* |
| **Innervation volume per taste cell** | 53.35±17.15 | 120.7±43.77 | p=0.0002* |
|  | **Mouse (n=8)** | **Human (n=9)** | **p-value** |
| **Number of synaptic contacts per taste cell** | 2.62±2.80 | 2.35±1.10 | p=0.7962 |
| **Mean volume of synaptic contact (μm<sup>3</sup>)** | 0.43±0.27 | 0.94±0.94 | p<0.0001* |

Standard deviations are shown. Differences with a p-value <0.05 highlighted in light gray.
\* Innervation density = (Innervation volume / Taste bud sectional volume) \* 100

Outliers for all measurements were detected using the robust outlier (ROUT) test with a maximum desired false discovery rate of 1%; this test was developed to identify outliers when fitting data with nonlinear regression (Motulsky and Brown 2006). The only outliers detected were in the data pertaining to individual synaptic contact size, in which 37 outliers (n=16 of 170 values for mice, n=21 of 164 values for humans). These outlier puncta were then manually examined with Imaris to determine if they comprised multiple individual synaptic contacts which have been merged as a single object, a confound of this analysis. Of the 37 points, 26 comprised multiple individual objects and were manually divided to permit inclusion as two individual objects; the remaining 11 were determined to be single objects, albeit large ones. After this processing, the data once again went through a ROUT analysis and 17 objects were identified as outliers (n=0 of 197 values for mice, n=17 of 182 values for humans). These were again visually examined, and none were considered to comprise multiple adjacent objects. We therefore considered these outliers as biological features in terms of determining the count and size of CALHM1 puncta. Accordingly, all objects were included in the final analysis including those identified as statistical outliers.

## Results

### Human taste buds are larger than mouse taste buds but contain similar numbers of taste cells

Human and mouse fungiform taste buds show similar structural and characteristic features including the presence of a taste pore, GNAT3 and PLCβ2-immunoreactive taste cells, and gustatory nerve fibers showing positive immunohistochemical staining for P2X3 (**Figure 1**). However, human and mouse fungiform taste buds are different sizes. Although they have similar widths (**Table 3**, **Figure 1**), human taste buds are 53% taller than mouse taste buds (67.7μm in humans vs. 43.1μm in mice). This feature is reflected in a 55% larger taste bud longitudinal sectional area (1901μm^2^ in humans vs. 2054μm^2^ in mice) defined as the taste bud area measured in the centermost image from the stack (**Figures 1, 2**). Accordingly, taste bud sectional volume – defined as the total taste bud volume calculated from the 6µM stack – is also larger in humans., Given the larger size of human taste buds, it might follow that human taste buds also contain higher numbers of DAPI-stained nuclear profiles of taste cells within the taste bud. However, mouse and human taste buds show similar nuclear profile counts although the nuclear profiles of human taste buds are slightly larger than those of mice (**Figure 2**). An Abercrombie correction to the taste cell count was used due to this difference in size between mouse and human taste cell nuclei. In both mice and humans, taste cell counts appear to increase proportionally to the taste bud sectional volume as has been previously noted in mice (Ohtubo and Yoshii 2011). Counts of PLCβ2-immunoreactive cells compared to the counts of nuclei suggest that approximately 26% of cells in taste buds in fungiform papillae in humans are immunoreactive for PLCβ2 compared to 30.6% in mouse as calculated from data in (Ohtubo and Yoshii 2011) (**Table 4**).

**Figure 1.**
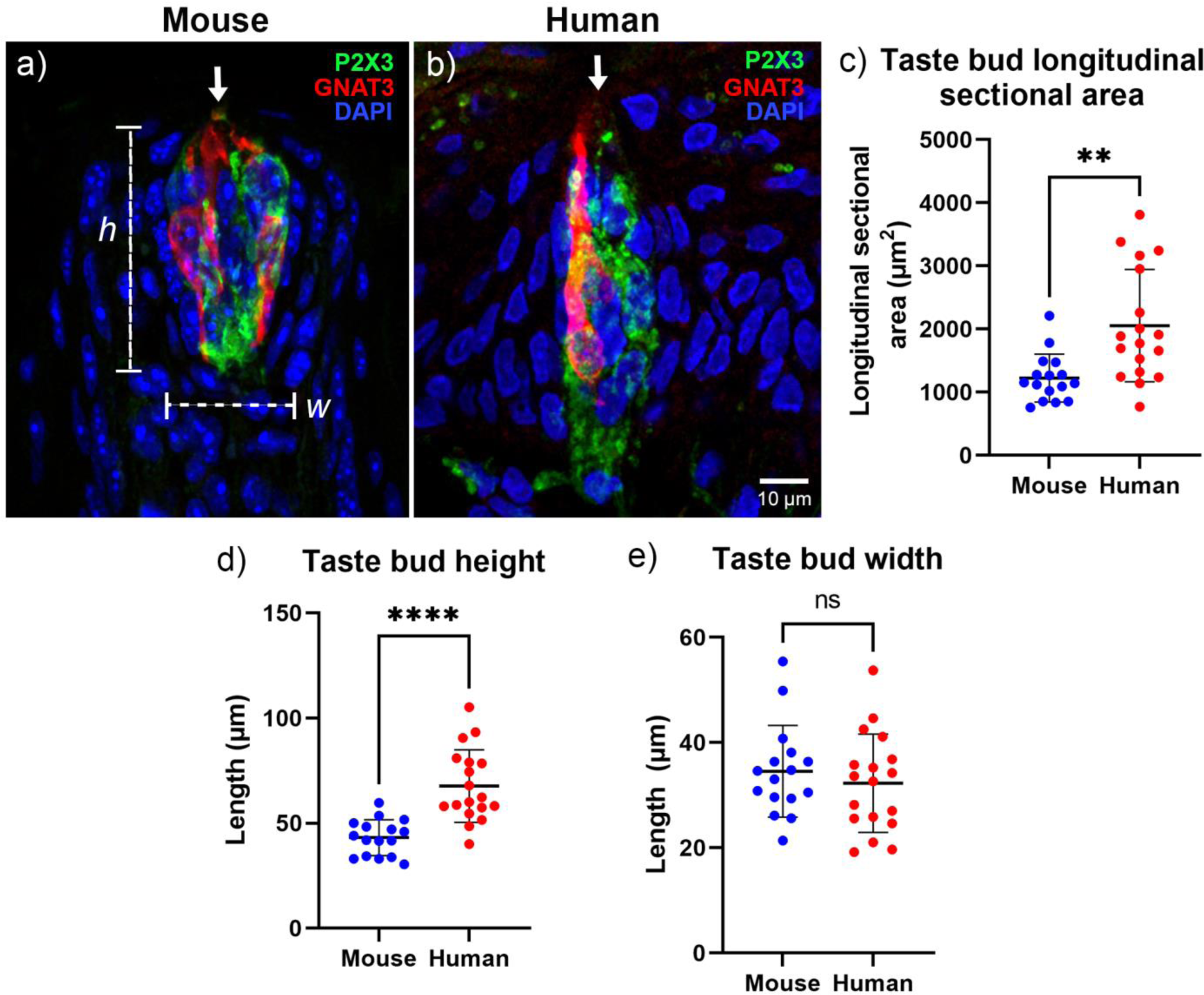
Taste buds have grossly similar structure in mouse (a) and human (b) fungiform papillae. White arrows indicate the taste pore. Gustatory nerve fibers (green) surround elongate taste cells (red). Human taste bud longitudinal sectional areas were measured and are significantly larger than mouse taste bud longitudinal sectional areas (d) (unpaired t-test, p=0.0014). Taste bud heights (d) and widths (e) were measured in human (n=18) and mouse (n=16) taste buds. Taste bud heights differ between human and mouse taste buds (unpaired t-test, p<0.0001) while taste bud widths do not (unpaired t-test, p=0.4723). Green = P2X3, a marker for gustatory nerve fibers. Red = GNAT3, a marker for a subset of type II taste cells. Blue = DAPI, a general nuclear stain.

**Figure 2.**
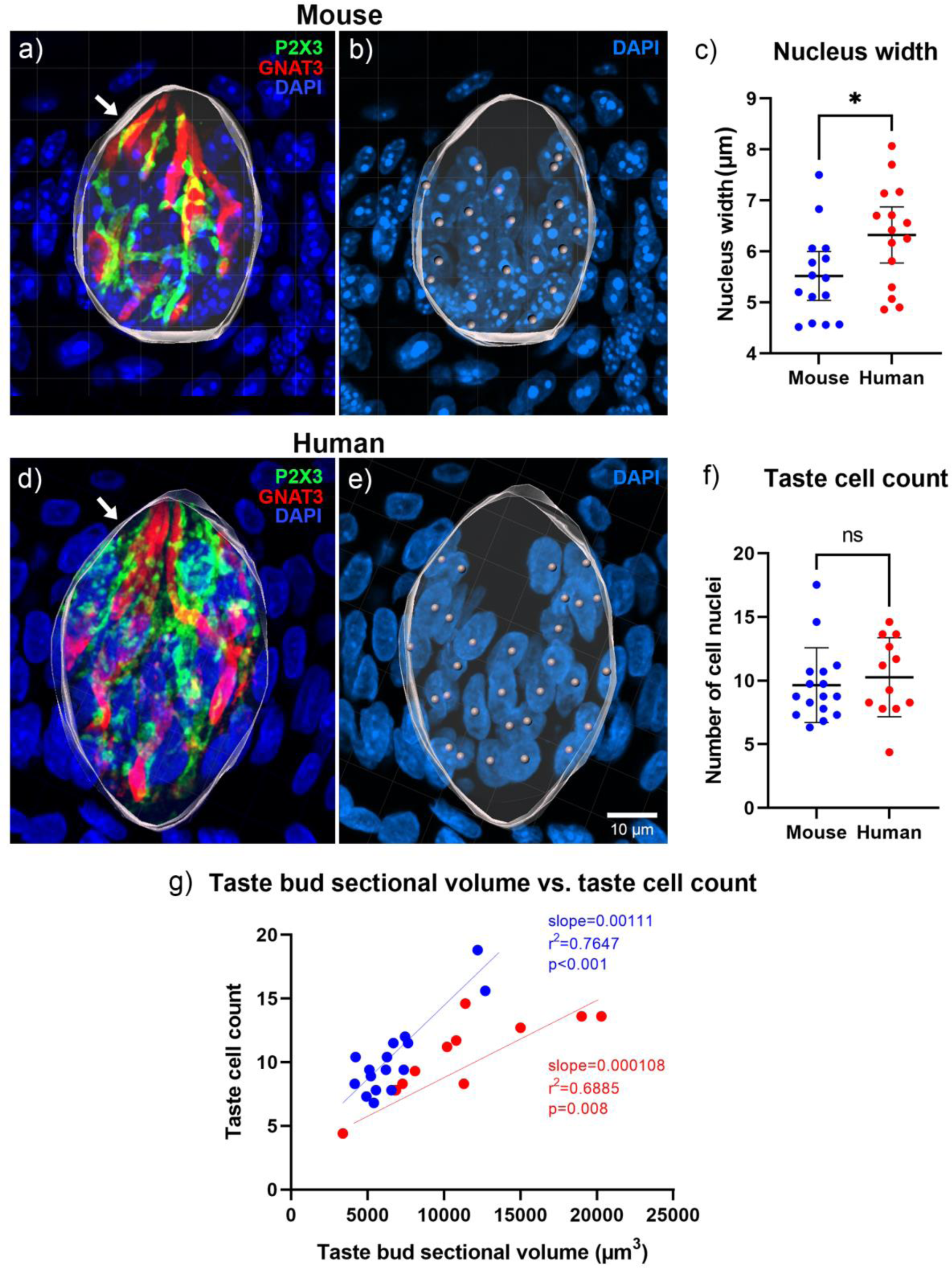
Mouse (a, b) and human (d, e) taste bud margins were outlined across a 6μM z-stack to create a three-dimensional taste bud sectional volume, indicated by the white arrows. Green = P2X3, a marker for gustatory nerve fibers. Red = GNAT3, a marker for a subset of type II taste cells. Blue = DAPI, a general nuclear stain. (c) Nuclear profile widths are significantly larger in humans (unpaired t-test, p=0.0256). (f) Abercrombie-corrected counts of mouse (b) and human (e) taste cell nuclear profiles reveal similar counts across both species (unpaired t-test, p=0.5945). (g) Nuclear profile counts compared to taste bud volume shows that generally, nuclear profile counts increase with increased taste bud volume in both species (linear regression). Blue, mouse; red, human.

**Table 4.** Counts and percentage of PLCβ2+ cells within human taste buds.

| **Subject** | **PLCβ2 count** | **Percentage of PLCβ2+ cells within the taste bud** |
| --- | --- | --- |
| 6 | 10 | 43.5% |
| 6 | 7 | 43.8% |
| 8 | 4 | 14.3% |
| 10 | 5 | 17.9% |
| 11 | 3 | 18.8% |
| 12 | 3 | 16.7% |
| 12 | 5 | 23.8% |
| 13 | 6 | 30.0% |
| ***Mean*** | **6.5±2.65** | **26.1%±0.1** |

### Human taste buds are more densely innervated than mouse taste buds

Both human and mouse fungiform taste buds show robust innervation by P2X3-immunoreactive nerve fibers (**Table 3**, **Figure 3**). In humans, however, these fibers are more plentiful and occupy 40% more volume within the 6μm z-stack compared to mice. Since human and mouse taste buds have similar numbers of taste cells, this finding means that human taste buds exhibit twice the innervation volume per taste cell profile, i.e., innervation density, compared to mouse taste buds (**Figure 3**). Observationally, nerve fibers within mouse fungiform taste buds also tend to occupy the lower portion of the taste bud more densely although individual fibers may extend towards the top of the taste bud. Human taste buds, however, show a dense plexus of nerve fibers throughout the height of the taste bud with fibers showing complex branching patterns. Finally, as in mice, larger taste buds in humans have more nerve fibers although innervation volume does not demonstrate a strong positive relationship with taste cell count in either mice or humans (**Figure 3**).

**Figure 3.**
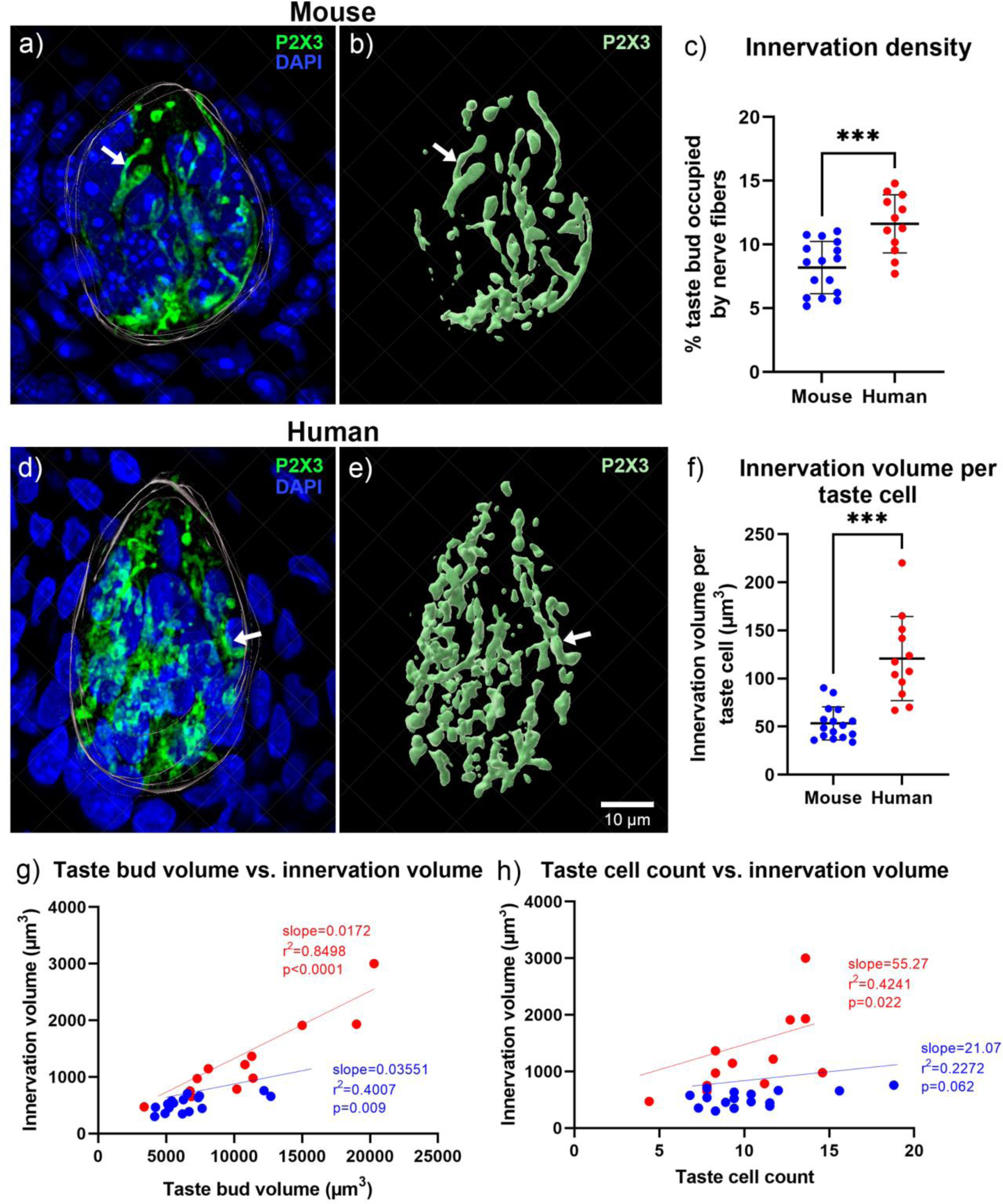
Mouse (a, b) and human (d, e) gustatory nerves were segmented within a 6μm z-stack to create a three-dimensional volume of gustatory innervation (b, e). Arrows indicate examples of corresponding areas in the raw image and in the three-dimensional reconstruction. (c) Innervation density is significantly larger in human than in mouse taste buds (unpaired t-test, p=0.0005). (f) Innervation volume per taste cell is higher in humans than in mice (unpaired t-test, p<0.0001). Innervation volume also appears to increase more in humans as taste bud volume increases (g) although increases in taste cell count do not appear to be strongly correlated with increased innervation volume (h) (linear regression, p-values on graph). Green = P2X3, a marker for gustatory nerve fibers. Blue = DAPI, a general nuclear stain.

### Channel-type synaptic contacts in human taste buds are larger than in mice

Human taste buds contain cells immunoreactive for PLCβ2, which has been used as a marker for type II taste cells in mice (**Figure 4**). In mice, PLCβ2 cells exhibit punctate immunoreactivity for CALHM1 and form appositional contacts with P2X3-immunoreactive nerve fibers characteristic of channel-type synapses (Ma *et al*. 2018; Romanov *et al*. 2018; Taruno *et al*. 2013). Human samples demonstrate punctate immunoreactivity at points of contact between PLCβ2-immunoreactive cells and P2X3-immunoreactive nerve fibers suggestive of channel-type synapses. Further, CALHM1 puncta are exclusively associated with PLCβ2 cells in human taste buds as in mice.

**Figure 4.**
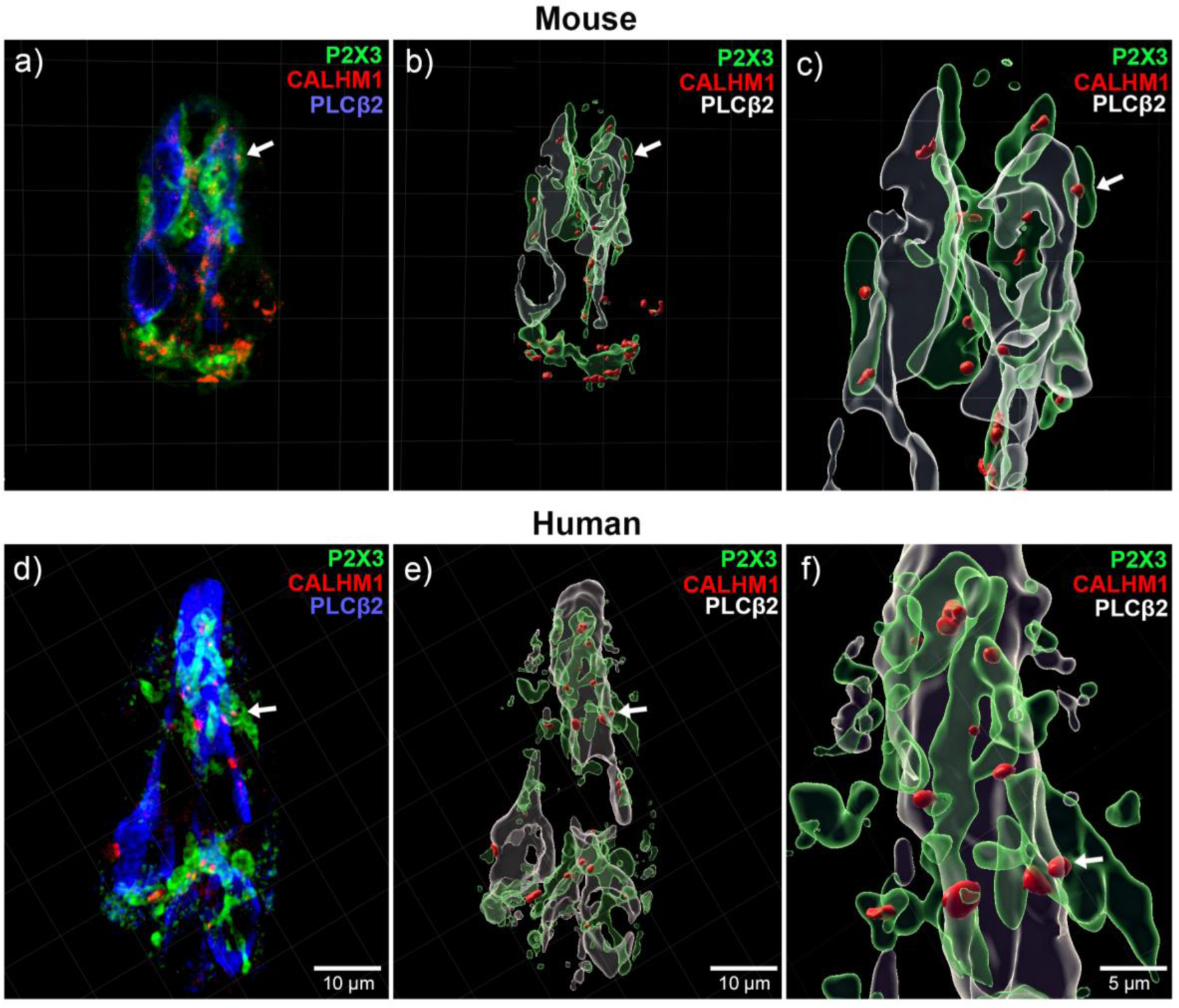
Human and mouse taste buds contain cells immunoreactive for PLCβ2, a marker of Type II cells in mice. In both species, these cells exhibit CALHM1 puncta situated at points of contact between the PLCβ2+ taste cells and gustatory nerve fibers. On average, the CALHM1+ puncta are larger in humans than in mice. Green = P2X3, a marker for gustatory nerve fibers. Red = CALHM1, a marker for synaptic contact. Blue = PLCβ2, a type II taste cell marker.

CALHM1 immunoreactivity was then used to measure the number and sizes of synaptic contacts in mouse and human fungiform taste buds (**Figure 5**). Although the number of synaptic contacts per 6μm section volume does not significantly differ between mouse and human taste buds, the average sizes of the synaptic contacts are twice as large in human fungiform taste buds (**Table 3**, **Figure 5**) with a substantial number of these contacts being larger than even the largest puncta in mouse taste buds. Finally, examination of the density of synaptic contact - whether calculated through innervation volume, total sectional volume, or taste cell count – does not show any significant differences between mouse and human taste buds (**Figure 5**).

**Figure 5.**
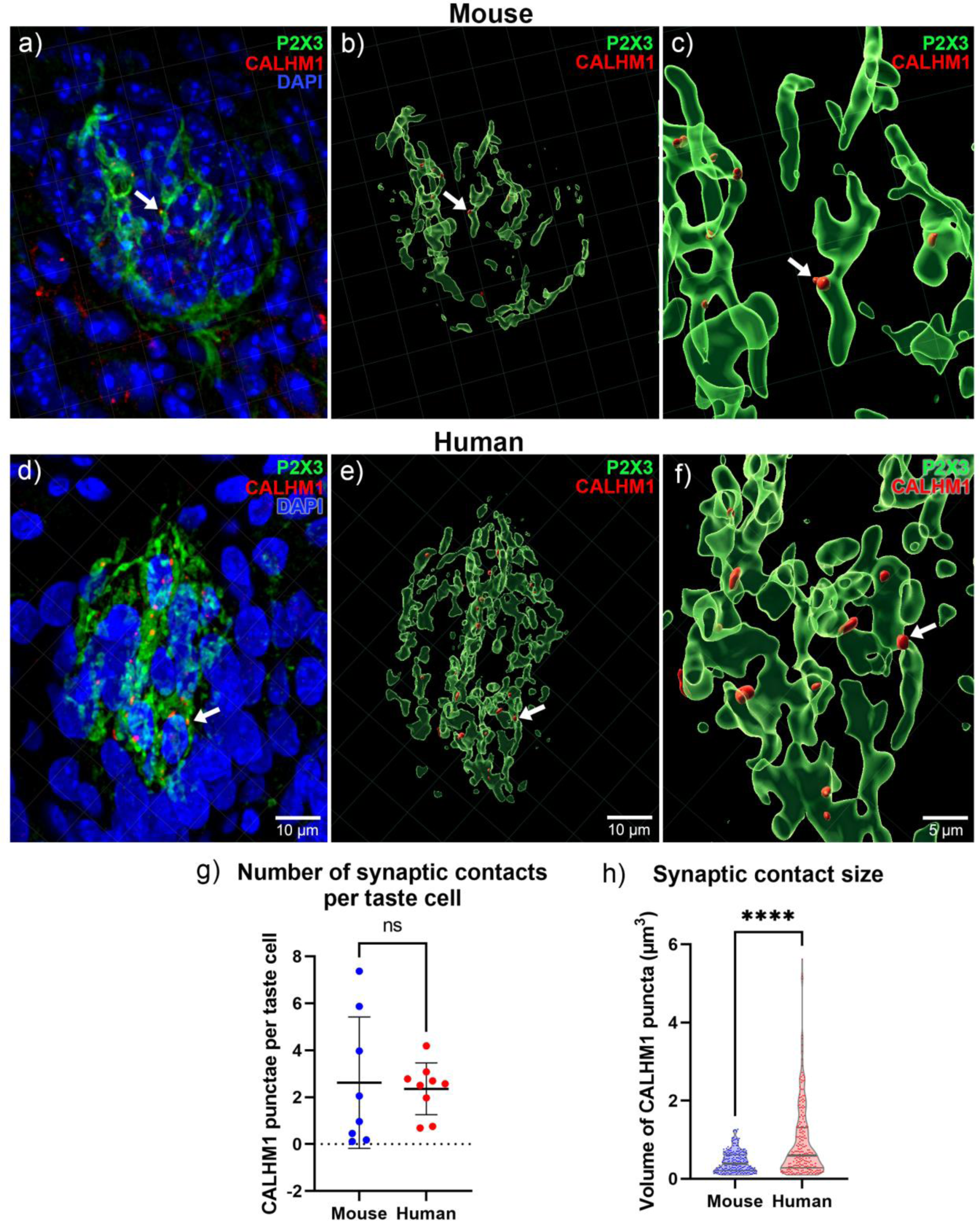
Channel-type synaptic contacts for mouse (a-c) and human (d-f) were measured within the 6μm image stack to generate counts and sizes of synaptic contact using CALHM1 as a marker. In both human and mouse taste buds synaptic contacts were defined as immunoreactive puncta situated within 0.2µM of P2X3-positive nerve fibers see Methods and Appendix for details). (g) The number of synaptic contacts per taste cell, does not significantly differ between species (unpaired t-test, p=0.80), but human synaptic contacts are significantly larger than mouse synaptic contacts, (mean size of a mouse synaptic contact=0.4335μm^3^ vs. a human synaptic contact=0.9365μm^3^) (c, f, h) (p<0.001). Green = P2X3, a marker for gustatory nerve fibers. Red = CALHM1, a marker for synaptic contact. Blue = DAPI, a general nuclear stain. n=8 taste buds for mouse, n=9 for human.

## Discussion

Human taste buds are generally similar in structure to those in mice but show significantly larger longitudinal sectional area (a measure of volume), increased innervation density, and larger synaptic contacts compared to those in mice. While a difference in sectional taste bud area may be expected given the significant size difference between a human and a mouse, the magnitude of this difference is nowhere near the 10-fold difference in overall size of the tongue between the two species.

Prior work reports that human and mouse taste buds have similar widths, but the increased height and longitudinal sectional area of human taste buds has not been previously examined (Miller 1989; Srur *et al*. 2010). It is possible that the taller taste buds of humans merely reflect a thicker tongue epithelium. Thus, to extend from basal lamina to the surface, a cell in a human taste bud must be longer than an equivalent cell in mouse. Notably, the number of taste cells is similar between mouse and human taste buds despite the increased height, which contributes to an overall larger taste bud volume in humans. However, taste cell nuclei are wider in humans than in mice, perhaps illustrating the possibility that in humans, the same number of taste cells fill in the larger volume of the taste bud. It is noteworthy that the number of cells is roughly similar between mouse and human, suggesting that this feature may be related to initial limitations on induction of the formation of taste buds from a limited number of progenitor cells in humans as well as mice (Hall *et al*. 1999; Liu *et al*. 2013; Stone *et al*. 2002; Thirumangalathu *et al*. 2009).

The higher innervation density in human taste buds is present despite a larger taste bud volume. This suggests that either more nerve fibers innervate each taste bud in humans, or that innervating fibers are more highly branched than in the murine system. Additionally, the nerve fibers in human taste buds appear to innervate apical portions of the taste bud more extensively than in mice **(c.f. Figure 3a,b with Figure 3d,e)**.

We show here for the first time that human taste buds, as in mice, contain the requisite components for channel synapses responsible for purinergic neurotransmission of type II taste cells, i.e., the presence of CALHM1 puncta within a PLCβ2-immunoreactive cell in apposition to P2X3-immunoreactive gustatory afferents (Taruno *et al*. 2013). The importance of purinergic signaling in taste function in humans is evidenced by taste dysfunction following treatments with purinergic receptor antagonists (McGarvey *et al*. 2022; Morice *et al*. 2021; Morice *et al*. 2019).

Although the number of synaptic contacts per taste cell is similar in mice and humans, synaptic contacts in human taste buds are, on average, twice as large as those in mice, although some channel synapse in humans appear considerably larger than the largest such contacts in mice. Presumably a larger channel synapse would permit greater flux of ATP through the activated channel. What this means in terms of transmission of activity in the system remains to be determined.

A confound in evaluating synaptic features was with the use of the CALHM1 antibody, which being generated in a mouse, yields higher non-specific labeling in the mouse tissue even with the use of an isotype-specific secondary antibody. Much of this background could be eliminated by using a lower limit particle size threshold, but even with this threshold, the synaptic puncta size in humans was still considerably larger than in mice. We tested several criteria to determine synaptic number and contact volume, i.e., thresholding images individually, changes in pre-processing steps, etc., but the results reached the same conclusion albeit with differing levels of variability. The results reported here use the same thresholding criteria across all images.

Altogether, these results suggest that baseline synaptic architecture, i.e., channel-type synapses formed between type II and PLCβ2-immunoreactive cells, is conserved between mice and humans. Further work is required to identify if other differences, particularly those related to molecular features of taste cell types, innervation, and synaptic structure, exist. Examining structural differences represents a rational comparative approach to uncovering where differences in taste signaling may lie between the two species. Such work might determine whether subsets of type II cells vary in CALHM1 synaptic contact number or size compared to mice, as variations in this could suggest functional differences in the detection and transmission of type II cell-mediated taste signals between the two species. Similarly, questions remain as well about the nature of these appositional contacts in humans and whether they also contain atypical mitochondria as in mice (Romanov *et al*. 2018). Finally, determining the relative numbers and ratios of type II to type III cells in human taste buds compared mouse could also be a clue towards uncovering whether functional differences might exist for type II or type III cell-mediated taste signaling between mammalian species.

## Data availability

The data underlying this article will be shared on reasonable request to the corresponding author.

## Acknowledgements

The authors thank Sue Kinnamon and Linda Barlow for comments and suggestions on this manuscript, Vijay Ramakrishnan for providing human fungiform tissues, Mei Li for assistance with immunohistochemical staining, and Dylan Volz for assistance with automating image preparation. The authors thank Adam Almeida, Katherine Given, and Eric Giedzinski for assistance with Imaris. The authors thank Wayne Rasband for development of ImageJ, (U. S. National Institutes of Health, Bethesda, Maryland, USA, https://imagej.nih.gov/ij/, 1997-2018) as well as Erich Giedzinski from Imaris for his help with developing the image analysis methods used here.

This work was supported by NIDCD DC019043-01A1 and R01DC014728.

## Supplementary methods

1. DAPI counting
  a. From the Object menu, choose the option to create a new Spots layer. Select the option to **Skip automatic creation and edit manually**
  b. Use **select mode** to manually place spots to count nuclei (DAPI) or cells (GNAT3). To switch from navigate mode to select mode quickly, press the escape key. Then, a spot can be placed anywhere on the image by holding down the shift key and left clicking.
  c. The cell count will be found in the **Tools** section under the **Statistics** tab.
2. Taste bud section volume
  a. Deselect the **Volume** option in the main **Object** menu.
  b. From the **Object** menu, choose the option to **Add new surfaces**. Then, select the option to **Skip automatic creation and edit manually**, and finally choose the **Contour** option.
  c. To trace the border of the taste bud, click on **Draw** and observe the arrow in the 3D view now appearing as a + symbol. Use the slicer tool to move through each optical section and outline the taste bud in each one.
  d. After outlining all the contours in each section, click on **Create Surface**. This surface was renamed to **Taste bud volume**. The volume object now appears in the main **Object** menu. The volume of the taste bud is in the **Tools** section under the **Statistics** tab.
3. Taste bud innervation volume
  a. Click the **pencil** icon in the **Tools** section for the **Taste bud volume** and select **Mask All**.
  b. Select the **P2X3** channel in the Channels dropdown menu. **Create Duplicate Channel** should already be checked. Type **0** in the box for **Set voxels outside surface to:**.
  c. A new channel will appear in the **Display Adjustment** window. This is an unaltered copy of the **P2X3** channel within the **Taste bud volume** only.
  d. From the **Object** menu, choose the option to **Add new surfaces**. Unselect the **Skip automatic creation, edit manually** option.
  e. Click through the blue arrows to observe the steps of thresholding. Select **Background Subtraction** and the **Diameter of largest Sphere which fits into the Object** should be set to 1.35. Click the green double arrow to finish surface creation, renamed to **P2X3**.
  f. The volume of innervation is in the **Tools** section under the **Statistics** tab.
4. CALHM1 volume and puncta
  a. Click the **pencil** icon in the **Tools** section for the **Taste bud volume** and select **Mask All**.
  b. Select the **CALHM1** channel in the Channels dropdown menu. **Create Duplicate Channel** should already be checked. Type **0** in the box for **Set voxels outside surface to:**
  c. A new channel will appear in the **Display Adjustment** window. This is an unaltered copy of the **CALHM1** channel within the **Taste bud volume** only.
  d. From the **Object** menu, choose the option to **Add new surfaces**.
  e. Click through the blue arrows and set thresholding steps in the following to:
  **i. Surface smoothing = 0.2 and background subtraction, largest sphere = 0.5.**
  **ii. Thresholding = 10.9**
  1. This thresholding step was chosen by individually thresholding each taste bud and then using the mean of these thresholding values to establish an average thresholding value to apply to all the CALHM1-stained taste buds.
  iii. Assign two filters to eliminate any detected puncta that are noise:
  **1. Shortest distance to P2X3 = 0.2**
  **2. Volume = 0.1**
  Click the green arrow to finish surface creation, renamed to **CALHM1**.
  The volume and number of CALHM1 puncta are in the **Tools** section under the **Statistics** tab.

## Notes

### Competing Interest Statement

The authors have declared no competing interest.

